# Chemical depletion of phagocytic immune cells reveals dual roles of mosquito hemocytes in *Anopheles gambiae* anti-*Plasmodium* immunity

**DOI:** 10.1101/422543

**Authors:** Hyeogsun Kwon, Ryan C. Smith

## Abstract

Mosquito innate immunity is comprised of both cellular and humoral factors that provide protection from invading pathogens. Immune cells, known as hemocytes, have been intricately associated with these immune responses through direct roles in phagocytosis and immune signaling. Recent studies have implicated hemocytes as integral determinants of anti-*Plasmodium* immunity, yet little is known regarding the specific mechanisms by which hemocytes limit malaria parasite survival. With limited genetic tools to enable their study, we employed a chemical-based treatment widely used for macrophage depletion in mammalian systems for the first time in an invertebrate organism. Upon its application in *Anopheles gambiae*, we observe distinct populations of phagocytic immune cells that are significantly depleted, causing high mortality following bacterial challenge and an increased intensity of malaria parasite infection. Through these studies, we demonstrate that phagocytes are required for mosquito complement recognition of invading ookinetes, as well as the production of prophenoloxidases that limit oocyst survival. Through these experiments, we also define specific sub-types of phagocytic immune cells in *An. gambiae*, providing new insights beyond the morphological characteristics that traditionally define mosquito hemocyte populations. Together, this study provides the first definitive insights into the dual roles of mosquito phagocytes in limiting malaria parasite survival, and illustrates the use of clodronate liposomes as an important advancement in the study of invertebrate immunity.

## Introduction

Innate immune defenses are essential for combating infectious pathogens throughout the animal kingdom (1). In insects, innate immunity is comprised of cellular and humoral components that coordinate killing responses through phagocytosis, melanization, and pathogen lysis. Immune cells, known as hemocytes, are integral to both cellular and humoral responses, respectively serving in direct or indirect roles to limit pathogen survival. Characterized largely by morphology, three circulating hemocyte subtypes have been described in mosquitoes with limited biochemical or genetic characterization (2–4). Functionally analogous to vertebrate macrophages, mosquito granulocytes are characterized by their phagocytic ability and adherence to tissues or foreign surfaces (3, 4). Oenocytoids have primarily been implicated in the production of phenoloxidases that lead to melanization responses (3–5), while prohemocytes are hypothesized to function as progenitor cells or as potentially smaller cells with phagocytic ability (3, 6).

Previous studies have demonstrated that *Anopheles gambiae* hemocytes respond to blood-feeding (7–10), as well as bacterial and malaria parasite challenge (3, 11), eliciting changes in their cellular profiles at the transcriptional (12, 13) and proteome levels (10). This has led to the identification of several mosquito hemocyte components with integral roles in phagocytosis (14, 15) and anti-*Plasmodium* immunity (10, 12, 16). In addition, hemocyte differentiation in response to malaria parasite challenge has been described as an integral requirement of mosquito immune priming (17–19) and the “late-phase” immune responses targeting *Plasmodium* oocyst survival (20–22). However, studies of mosquito immune cell function have been severely limited by the lack of genetic tools and resources to better understand the contributions of individual hemocyte sub-types to mosquito innate immune function.

In an effort to overcome these limitations, we have employed clodronate liposomes (CLD) to chemically deplete phagocytic cell populations in *Anopheles gambiae* using techniques that have been widely established in mammalian systems to ablate macrophage populations (23–25). Through this methodology, we take advantage of the phagocytic properties of mosquito immune cells to better understand their functional contributions to pathogen challenge and anti-*Plasmodium* immunity. Following CLD treatment, phagocyte depletion was confirmed using multiple methods of validation (light microscopy, flow cytometry, qRT-PCR, and immunofluorescence assays) which conferred significant impacts on mosquito survival after bacterial challenge. In addition, phagocyte depletion notably impaired *Plasmodium* killing responses at both the ookinete and oocyst stages, providing new mechanistic understanding into the immune components the influence parasite survival in the mosquito host. Together, these data represent a significant advancement in the genetic techniques used to study invertebrate immune cell function, as well as define new integral roles of phagocytic immune cells in mosquito anti-*Plasmodium* immunity.

## Materials and Methods

### Mosquito rearing

*An. gambiae* Keele strain mosquitoes (26, 27) were reared at 27°C and 80% relative humidity, with a 14/10 hour day/night cycle. Larvae were fed on fish flakes (Tetramin, Tetra), while adult mosquitoes were maintained on 10% sucrose solution.

### *Plasmodium* infection

Female Swiss Webster mice were infected with *P. berghei*-mCherry strain as described previously (20, 21). Infected mosquitoes were maintained at 19°C until individual mosquito midguts were dissected to count oocyst numbers by fluorescence microscopy (Nikon Eclipse 50i, Nikon).

### Phagocyte depletion using clodronate liposomes

Naïve female mosquitoes (4- to 6-day old) were injected intra-thoracically with either 69 nl of control liposomes (LP) or clodronate liposomes (CLD) (Standard macrophage depletion kit, Encapsula NanoSciences LLC) using a Nanoject II injector (Drummond Scientific). Initial experiments were performed using different dilutions of the stock concentrations using LP and CLD in 1×PBS (1, 1:2, 1:5, and PBS only) to determine CLD efficacy on phagocyte depletion and its effects on mosquito survival. All subsequent experiments were performed using a 1:5 dilution of control or clodronate liposomes in 1×PBS. Liposome injections were performed either on naïve mosquitoes one day prior to blood feeding or *P. berghei* challenge (pre-treatment), or on mosquitoes 24 h after *P. berghei* infection (post-treatment).

### Hemolymph perfusion and hemocyte counting

Hemolymph perfusion and hemocyte counting were performed as previously described (20, 21). Hemolymph was collected from pre-treated mosquitoes at 24 h (24 h naïve), 48 h naïve, 24 h blood-fed (24 h BF), 24 h *P. berghei* infection (24 h *P.b*) and post-treated mosquitoes at 48 h *P. berghei* infection (48 h *P.b*) using anticoagulant solution (vol/vol, 60% Schneider’s insect medium, 10% fetal bovine serum and 30% citrate buffer; 98 mM NaOH, 186 mM NaCl, 1.7 mM EDTA, and 41 mM citric acid, pH 4.5) as described previously (20, 21). Hemolymph (10 μl) from an individual mosquito was perfused through an incision made in the lateral abdomen. Hemocytes were quantified counting ~200 cells per individual mosquito, and the hemocyte subtypes were evaluated by morphological differences using a disposable Neubauer hemocytometer slide (C-Chip DHC-N01, INCYTO).

### *In vivo* hemocyte staining using CM-DiI

To visualize the depletion of phagocytes following CLD treatment, hemocytes were stained as previously (6, 11, 20, 28) using CM-DiI (Vybrant CM-DiI, Life Technologies) and FITC-conjugated wheat germ agglutinin (WGA, Sigma). Briefly, mosquitoes pre-treated with either control liposomes or clodronate liposomes were challenged with *P. berghei*, then ~24 hrs post-infection were injected with 138 nl of 100 μM CM-DiI and incubated for 20 min at 19°C. Perfused hemolymph (~10 μl) was collected onto a multitest glass slide (MP Biomedicals) and hemocytes were allowed to adhere to the slide for 30 min. Without washing, 4% paraformaldehyde was added to each well for fixation. After incubating at RT for 30 min, cells were washed 3 times in 1× PBS for 5 min, then blocked with 1% BSA in 1xPBS for 30 min before incubating with WGA (1:125) overnight at 4°C. Prior to visualization, slides were washed three times in 1× PBS for 5 min, then mounted with ProLong^®^Diamond Antifade mountant with DAPI (Life Technologies).

### Flow cytometry

Flow cytometry analyses was performed to confirm phagocyte depletion following CLD treatment. Mosquitoes pre-treated with CLD under naïve, 24 h blood-fed, or 24 h *P. berghei*-infected conditions were injected with red fluorescent FluoSpheres (1 μm, Molecular Probes) at a final concentration of 2 % (vol/vol) and allowed to recover for 2 hours at 19°C before perfusion. Hemolymph was collected from ~60 individual mosquitoes for each experimental treatment as described above using anticoagulant solution as previously described (20, 21). Perfused cell samples were fixed in 4% paraformaldehyde for 1 hour at 4°C, then centrifuged for 5 min at 2000 × g to pellet cells, while discarding the supernatant. Cells were washed two times in 1× PBS with an additional centrifugation step of 5 min at 2000 × g between washing steps. Samples were incubated with WGA (1:5000) and DRAQ5 (1:1000, Thermo Fisher Scientific) overnight at 4°C. Following incubation, cells were washed two times in 1× PBS to remove excess stain and then run on a BD FACSCanto cytometer (BD Biosciences). Data were analyzed with FlowJo software (FlowJo, LLC) using strict threshold values for gating as determined by a fluorescent bead-only sample to exclude events by size (forward scatter, FSC) and the use of unstained cells to determine cutoffs for positive WGA and DRAQ5 signals to remove any auto-fluorescence background. To measure the effects of CLD depletion, the percentage of phagocytic cells in LP and CLD samples were used for comparison. Flow cytometry experiments were performed three or more times from independent biological experiments for naïve, blood-fed, and *P. berghei*-infected conditions.

### RNA isolation and gene expression analyses

Total RNA was isolated from dissected mosquito tissues or from whole mosquito samples to examine gene expression using TRIzol (Thermo Fisher Scientific). 2 μg of total RNA was used a template for cDNA synthesis using the RevertAid First Strand cDNA Synthesis kit (Thermo Fisher Scientific). qRT-PCR was performed using PowerUp^™^SYBR^®^Green Master Mix (Thermo Fisher Scientific) with the ribosomal S7 protein transcript serving as an internal reference as previously (21). cDNA (1:5 dilution) amplification was performed with 250 nM of each specific primer pair using the following cycling conditions: 95°C for 10 min, 40 cycles with 95°C for 15 s and 65°C for 60 s. A comparative C_T_ (2^−ΔΔCt^) method was employed to evaluate relative transcript abundance for each transcript (29). A list of primers used for gene expression analyses are listed in Table S1.

### Phagocytosis assays

Phagocytosis assays were performed by injecting 69 nl of red fluorescent FluoSpheres at final concentration of 2 % (vol/vol) into naïve, 24 h blood-fed, or 24 h *P. berghei*-infected mosquitoes as previously (20). Following injection, mosquitoes were kept at 19°C for 2 h before hemolymph was perfused onto a multitest glass slide. Hemocytes were allowed to attach the slide for 30 min at room temperature (RT) and were then fixed with 4% paraformaldehyde for 30 min. Slides were washed 3 times in 1× PBS for 5 min each wash, then blocked in 1% of bovine serum albumin (BSA) for 30 min. To visualize cells, samples were incubated with 1:500 WGA overnight at 4°C. Hemocytes were washed three times in 1× PBS, then mounted with ProLong^®^Diamond Antifade mountant with DAPI. Hemocytes labelled with WGA and harboring red fluorescent beads were considered as phagocytes. Phagocytic activity was evaluated as the number of immune cells engulfing one or more fluorescent beads divided by total immune cell population (n=50/mosquito) counted from random chosen fields under a fluorescent microscope. The average number of fluorescent beads per phagocyte was also determined and is referred to as the phagocytic index.

### Bacterial challenge experiments

Bacteria were grown in LB media overnight at 37°C with 210 rpm. Bacterial cultures were centrifuged at 8000 rpm for 5 min, washed twice with 1× PBS, and resuspended in 1×PBS. ~24 h after pre-treatment with control or clodronate liposomes, mosquitoes (n=30) were injected with 69 nl of bacterial suspensions (100X dilution of *Serratia marcescens* or *Staphylococcus aureus* at OD_600_=0.4) using a nanoinjector. Mosquitoes pre-treated with control liposomes were also injected with 69 nl of 1× PBS to serve as an additional control. Following challenge, mosquitoes were maintained at 27°C and 80% relative humidity with mosquito survival monitored every 24 h for 10 days.

### Western blot analysis

Following liposome treatments, hemolymph was perfused from individual mosquitoes (n=15) at naïve or 24 h *P. berghei*-infected mosquitoes using incomplete buffer (anticoagulant solution without fetal bovine serum) containing a protease inhibitor cocktail (Sigma) (10). Hemolymph protein concentrations were measured using Quick Start^™^Bradford Dye reagent (Bio-Rad). Protein samples (~2 μg) were mixed with Bolt^™^LDS sampling buffer and sample reducing agent (Life Technologies), and heated at 70°C for 5 min before separated on 4-12% Bis-Tris Plus ready gel (Thermo Fisher Scientific). Samples were resolved using Bolt^™^MES SDS running buffer (Thermo Fisher Scientific) for 90 min at 100 V. Proteins were transferred to PVDF membrane in Bolt^™^Transfer buffer (Life Technologies) for 1 h at 20 V, and then blocked in TBST buffer (10 mM Tris base, 140 mM NaCl, 0.05% Tween 20, pH 7.6) containing 5% non-fat milk for 1 hour at RT. For western blotting, the membrane was incubated with a 1:1000 dilution of rabbit anti-TEP1 (30), rabbit anti-PPO6, or rabbit anti-serpin3 (SRPN3) antibodies (31) in TBST blocking buffer overnight at 4°C. Membranes were washed three times for 5 min in TBST, then incubated with a secondary anti-rabbit alkaline phosphatase-conjugated antibody (1:7500, Thermo Fisher Scientific) for 2 h at RT. Following washing in TBST, the membrane was incubated with 1-Step^™^NBT/BCIP (Thermo Fisher Scientific) to enable colorimetric detection. For comparative analysis between samples, densitometric analysis of protein bands was performed using ImageJ (https://imagej.nih.gov/ij/).

### TEP1 immunofluorescence assays

Analysis of TEP1 binding to invading ookinetes was performed by immunofluorescence similar to previous experiments (28, 30, 32, 33). Mosquitoes pre-treated with liposomes and infected with *P. berghei* were dissected at ~22-24 h post-infection. Midguts were dissected and briefly fixed in 4% paraformaldehyde for 40s, then the blood bolus was removed and the midguts briefly washed in 1× PBS before fixation in 4% PFA for 1 hour at RT. Following washing three times in 1× PBS, midguts were blocked in blocking buffer (1% bovine serum albumin and 0.1% Triton X-100 in 1xPBS) overnight at 4°C. Midgut sheets were incubated with mouse α-Pbs21 (1:500) and rabbit-TEP1 (1:500) primary antibodies in blocking buffer overnight at 4°C. After washing in 1×PBS, midguts were incubated with Alexa Fluor 568 goat anti-mouse IgG (1:500, Thermo Fisher Scientific) and Alexa Fluor 488 goat anti-rabbit IgG (1:500, Thermo Fisher Scientific) secondary antibodies in blocking buffer for 2 h at RT. Midguts were washed three times in 1×PBS, then mounted with ProLong^®^Diamond Antifade mountant with DAPI.

### RNA-Seq and differential gene expression analysis

Adult female mosquitoes were pre-treated with either control or clodronate liposomes as described above, then challenged with *P. berghei.* Approximately 24 h post-infection, mosquitoes were dissected in 1× PBS to dissociate the gut and reproductive organs from the abdominal wall. Total RNA was extracted from dissected abdomen (n=15) tissues using TRIzol (Thermo Fisher Scientific) for each treatment. The isolated RNA was further purified with the RNA Clean & Concentrator-5 kit (Zymo research) and quantified using a Nanodrop spectrophotometer (Thermo Fisher Scientific). RNA quality and integrity were measured using an Agilent 2100 Bioanalyzer Nano Chip (Agilent Technologies), and 200 ng of total RNA from four independent biological replicates was used to perform RNA-seq analysis. Libraries were prepared by the Iowa State University DNA Facility using the TruSeq Stranded mRNA Sample Prep Kit (Illumina) using dual indexing according to the manufacturer’s instructions. The size, quality, and concentration of the libraries was measured using an Agilent 2100 Bioanalyzer and a Qubit 4 Fluorometer (Invitrogen), then diluted to 2 nM based on the size and concentration of the stock libraries. Clustering of the libraries into a single lane of the flow cell was performed with an Illumina cBot. 150 base paired end sequencing was performed on an Illumina HiSeq 3000 using standard protocols.

Raw sequencing data was analyzed by the Iowa State genome Informatics Facility. Sequence quality was assessed using FastQC (v 0.11.5) (34), then paired end reads were mapped to the *Anopheles gambiae* PEST reference genome (AgamP4.9) downloaded from VectorBase (35) using STAR aligner (v 2.5.2b) (36). Genome indexing was performed using the genomeGenerate option and corresponding GTF file downloaded from VectorBase (version 4.7) followed by mapping using the alignReads option. Output SAM files were sorted and converted to BAM format using SAMTools (v 1.3.1) (37), and counts for each gene feature were determined from these alignment files using featureCounts (v 1.5.1) (38). Reads that were multi-mapped, chimeric, or fragments with missing ends were excluded. Counts for each sample were merged using AWK script and differential gene expression analyses was performed using edgeR (39). Differentially expressed genes with a q-score ≤0.1 were considered significant and were used for downstream analyses. Gene expression data have been deposited in NCBI’s Gene Expression Omnibus (40) and are accessible through GEO Series accession number GSE116156 (https://www.ncbi.nlm.nih.gov/geo/query/acc.cgi?acc=GSE116156).

Several candidate genes identified in our RNA-seq expression analysis were selected and measured by qRT-PCR to further validate the results of our gene expression experiments. Transcripts influenced by phagocyte depletion were selected according to significant fold-change values. Independent mosquito carcass samples ~24 h post-*P. berghei* infection were prepared from pre-treated liposome mosquitoes samples and were used for validation experiments. Total RNA isolation, cDNA synthesis and qRT-PCR were performed as described above. These same samples were used for additional follow-up experiments examining PPO gene expression. In addition, relative PPO gene expression was determined in total hemocyte samples collected from perfused hemolymph (n=60) following LP and CLD treatment in infected (~24 h) *P. berghei* mosquitoes. RNA was isolated using TRIzol and was then further purified with RNA Clean & Concentrator-5 kit. 200ng of total RNA (200 ng) was used for cDNA synthesis. Primers used are listed in Table S1.

### Gene silencing by dsRNA

RNAi experiments were performed with selected genes: *PPO2* (AGAP006258), *PPO3* (AGAP004975), *PPO4* (AGAP004981), *PPO5* (AGAP012616), *PPO6* (AGAP004977), *PPO9* (AGAP004978), *CLIPD1* (AGAP002422) and a putative leucine-rich immunomodulatory (LRIM) protein (AGAP001470). T7 primers were designed using the E-RNAi web application (http://www.dkfz.de/signaling/e-rnai3/idseq.php) and listed in Table S2. T7 templates for dsRNA synthesis were amplified from cDNA prepared from whole mosquitoes ~24 h post-*P. berghei* infection. PCR amplicons were purified using the DNA Clean & Concentration kit (Zymo Research). dsRNAs were synthesized using the MEGAscript RNAi kit (Life Technologies) according to the manufacturer’s instructions, and then resuspended in nuclease free water to 3 μg/μl after ethanol precipitation. Three to four day old mosquitoes were cold anesthetized and injected in the thorax with 69 nl (~200 ng) of dsRNA per mosquito. The effects of gene silencing were measured 2 days post-injection in whole mosquitoes (n=15) by qRT-PCR as described above. Although dsRNA targeting each individual PPO target gene were prepared following E-RNAi design (41), potential off-target effects on other PPO gene family members were examined to determine if the knockdown of a specific PPO dsRNA influences the expression of other PPO transcripts. Primers used for gene silencing and qRT-PCR experiments are listed in Table S1. To evaluate the effects of gene-silencing on malaria parasite infection, mosquitoes were challenged with *P. berghei* 2 days post-injection of dsRNA. Oocyst numbers were examined at either 2 days or 8 days post-infection.

### Phenoloxidase (PO) assay

Phenoloxidse (PO) activity was measured in pools of perfused hemolymph from pre-treated mosquitoes (n=15, 10 μl per mosquito) under naïve conditions, 24 h post-blood feeding, and 24 h post-*P. berghei* infection. Following perfusion, 10 μl of the total perfused hemolymph was mixed with 90 μl 3, 4-Dihydroxy-L-phenylalanine (L-DOPA, 4 mg/ml) dissolved in nuclease free water as previously described (42). PO activity was measured at 490 nm every 5 min for 30 min, then the final activity was measured at 60 min using a microplate reader (Molecular Devices).

### Hemocyte immunofluorescence assays

Hemocyte immunofluorescence assays were performed using a PPO6 antibody as previously described (8, 9). Mosquitoes pre-treated with either control liposomes or clodronate liposomes were injected with red fluorescent FluoSpheres at a final concentration of 2 % (vol/vol) approximately 24 h after *P. berghei* infection. Following incubation for 2 h at 19°C, hemocytes were perfused on a multi-well glass slide and allowed to adhere at RT for 30 min. Cells were fixed with 4% paraformaldehyde for 30 minutes, then washed three times in 1× PBS. Samples were incubated with blocking buffer (0.1% Triton X-100, 1% BSA in PBS) for 1 h at RT and incubated with rabbit-anti PPO6 (1:500) in blocking buffer overnight at 4°C. After washing 3 times in 1× PBS, an Alexa Fluor 488 goat anti-rabbit IgG (1:500) secondary antibody was added in blocking buffer for 2 h at RT. Slides were rinsed three times in 1× PBS and mounted with ProLong^®^Diamond Antifade mountant with DAPI. Hemocytes were screened for the presence of PPO6 signal and phagocytic activity. The percentage of PPO6^+^ cells displaying phagocytic ability were counted as a proportion of the total cell population. Approximately 200 randomly selected hemocytes were evaluated in liposome controls, while the entire cell population was counted in CLD-treated samples with less than 200 cells. Discernable differences in phagocyte populations between control and CLD-treated samples were determined based on morphology.

## Results

### Phagocyte depletion using clodronate liposomes

In an effort to understand the principal roles of phagocytic immune cells in mosquito immunity, clodronate liposomes were employed to chemically deplete phagocytes in *An. gambiae* (Fig. S1A). Widely used in studies of mammalian immunity (23–25), clodronate liposomes were first injected into adult female mosquitoes to titer the required concentration of liposomes needed for depletion. From these experiments, a 1:5 dilution of clodronate liposomes in 1xPBS was chosen for its efficacy in ablating phagocytes without negative impacts on mosquito survival (Fig. S1B). To determine the efficacy of cell depletion, granulocyte populations were evaluated from naïve, blood-fed, or *P. berghei*-infected mosquitoes based on morphology (Fig. S2) using a hemocytometer as previously (20, 43). Clodronate treatment reduced the percentage of granulocytes by ~40% in naïve mosquitoes when examined at either 24 or 48 hours (Fig. S2), yet higher levels of depletion (~90%) were observed in blood-fed or *P. berghei*-infected samples (Fig. S2). This increase in phagocyte depletion may be attributed to the enhanced phagocytic ability and capacity following the physiological changes that accompany blood-feeding (Fig. S3), presumably enabling a higher dosage of clodronate to the cell. Depletion was further validated by immunofluorescence of fixed hemocyte populations stained with DiI and WGA, which have previously been used to label hemocyte populations (11, 20, 28), demonstrating that immune cell populations were significantly reduced in CLD-treated mosquitoes when compared to liposome controls (Fig. 1A). This morphological data supports that granulocyte populations are significantly reduced following clodronate treatment.

**Figure 1.**
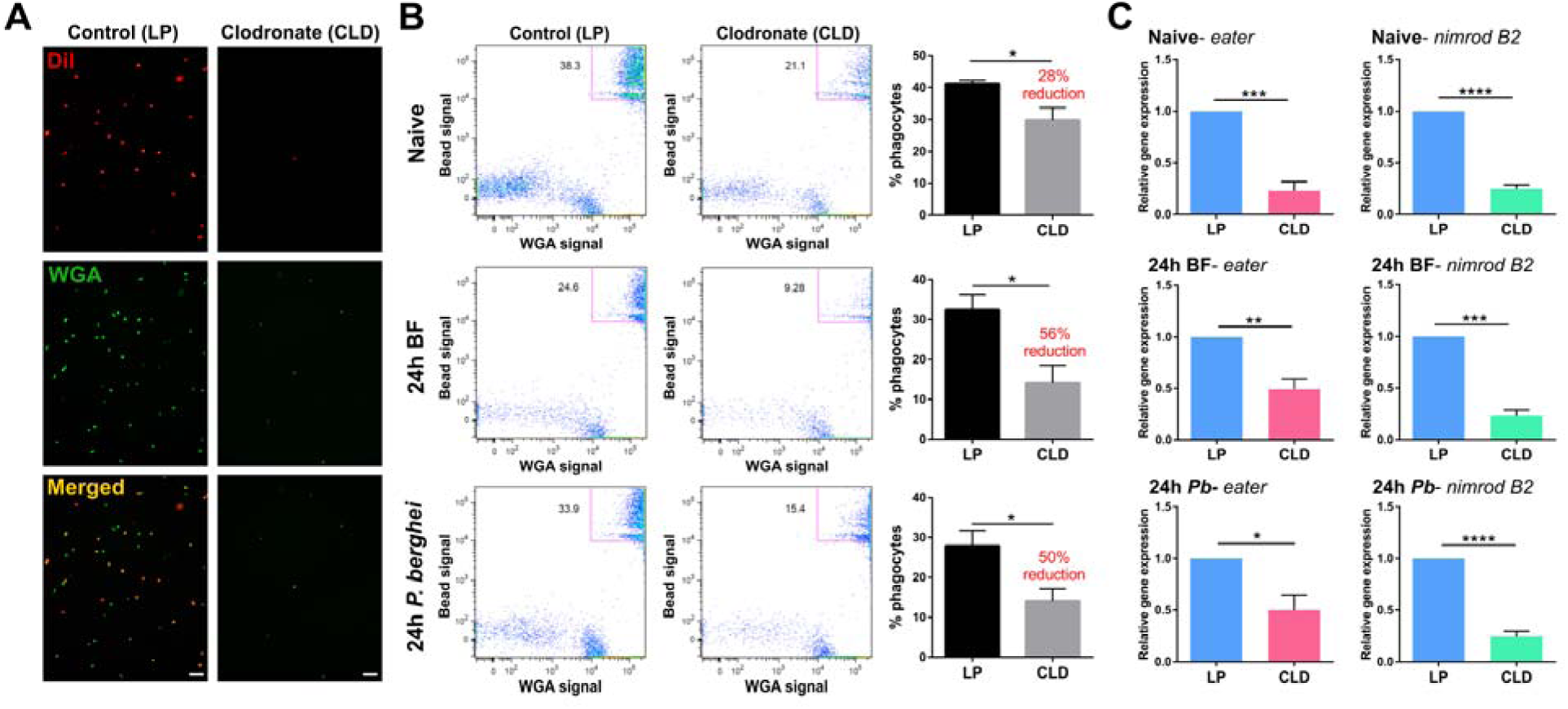
Mosquito phagocytes are significantly depleted following clodronate liposome (CLD) treatment. Following the injection of control (LP) or clodronate liposomes (CLD), mosquitoes were challenged with *P. berghei* and hemocytes were perfused 24 h post-infection. Hemocytes were stained with the hemocyte-specific markers, DiI (red) and WGA (green), to visualize the effects of CLD-treatment on hemocyte populations (**A**). Further validation was performed using flow cytometry to confirm the depletion of phagocytic hemocyte populations under naïve, 24 h blood-fed (24 h BF), and 24 h *P. berghei*-infected (24 h *P.b*) conditions (**B**). Representative flow cytometry experiments display the depletion of phagocytic hemocytes (as determined by the uptake of fluorescent beads and WGA staining), with bar graphs depicting a significant decrease in phagocytes following CLD treatment from three independent experiments (**B**). Blood-feeding, independent of pathogen challenge, caused phagocytes to be more susceptible to CLD treatment (**B**). Additional experiments with molecular markers of phagocytic cells, *eater* and *nimrod B2*, were used to further evaluate phagocyte depletion. Relative transcript levels of *eater* and *nimrod B2* expression were significantly reduced in CLD treated mosquitoes (**C**). Bars represent mean ± SEM of three independent replicates. Data were analyzed by unpaired t-test using GraphPad Prism 6.0. Asterisks denote significance (^⋆^*P* < 0.05, ^⋆⋆^*P* < 0.01, ^⋆⋆⋆^*P* < 0.001, ^⋆⋆⋆⋆^*p* < 0.0001). Scale bar: 20 μm.

Based on our previous work isolating phagocytic immune cells by their phagocytic properties (10), we employed a similar approach to distinguish the effects of clodronate treatment on phagocytic cells utilizing flow cytometry analyses. Using fluorescent beads to distinguish phagocytic cells from other hemocyte subtypes (Fig. S3; (44)), phagocytic cell populations could effectively be measured and compared between treatments using strict size cutoffs and signal thresholds (Fig. S4). Flow cytometry analyses revealed that CLD treatment significantly depleted mosquito phagocyte populations under naïve, blood-fed, and *P. berghei*-infected conditions (Fig. 1B; Fig. S5). Similar to our results with light microscopy (Fig. S2), the efficacy of phagocyte depletion is increased following a blood meal, independent of infection status (Fig. 1B). Additional validation of phagocyte depletion was performed by examining the transcripts of two well-characterized genes associated with hemocyte phagocytic function, *eater* and *nimrod B2* (45–48), by qRT-PCR. Under each experimental condition, the relative transcript abundance of *eater* and *nimrod B2* in CLD treated mosquitoes was significantly reduced when compared to control liposome (Fig. 1B). Together, these data provide strong support for the chemical depletion of mosquito phagocytic immune cells using clodronate liposomes.

In addition, our flow cytometry data argue that there are at least two distinct populations of phagocytic immune cells in *An. gambiae.* Under each experimental condition, phagocytic cells were noticeably segregated into upper and lower phagocyte populations (Fig.1B; Fig. S6A). Across physiological conditions, the upper population was more susceptible to CLD treatment, displaying significant reduction in the percentage of phagocytic cells. This is in contrast to the lower phagocyte population, which displayed little response to CLD treatment (Fig. S6A). Further quantification of these upper and lower populations by size (FSC) and granularity (SSC) demonstrate size differences under naïve conditions and significant disparities in granularity across feeding status (Fig. S6B). Similar comparisons between control and CLD treatments display no major differences between these upper and lower populations (Fig. S6C). These data suggest that these two populations are distinct cell types with different phagocytic potential, which may explain why the increased phagocytic activity in the upper cell population results in a stronger depletion following CLD treatment.

### Phagocyte depletion increases susceptibility to bacterial infections

To determine the influence of phagocyte depletion on immune function and host survival, control and CLD liposome treated mosquitoes were challenged with bacteria. Injury alone had little effect on mosquito survival, while challenging with either gram (+) or gram (-) bacteria had notable impacts on survivorship (Fig. 2). However, the impact of phagocyte depletion significantly decreased mosquito survival to *S. marcescens* (Fig. 2A) and S. *aureus* challenge (Fig. 2B). Mosquitoes were highly susceptible to *S. marcescens*, killing all CLD treated mosquitoes within 2 days post-challenge (Fig. 2A). Given the importance of phagocytic cells as immune sentinels required to remove invading pathogens (46, 48, 49), these experiments further demonstrate the effects of clodronate depletion and the important contributions of phagocytes as critical effectors of mosquito cellular immunity.

**Figure 2.**
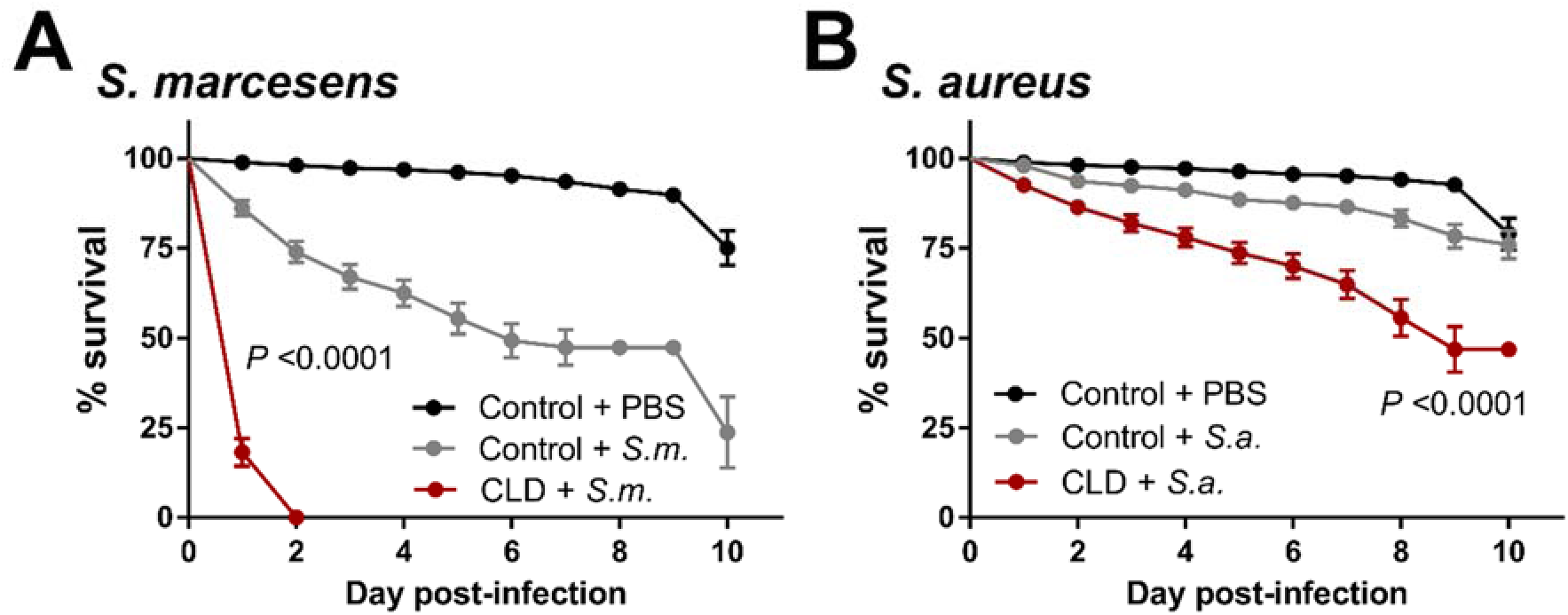
Depletion of phagocytic cells influences mosquito survival after bacterial challenge. Mosquitoes were treated with either control or clodronate liposomes, then subjected to injury (sterile PBS injection) or bacterial challenge. Survivorship was monitored in mosquitoes every day for ten days to evaluate the effects of *S. marcescens* (**A**) or *S. aureus* (**B**) challenge. Phagocyte depletion (CLD) in mosquitoes results in a high susceptibility to bacterial infections. Error bars represent the mean ± SEM of three independent replicates. In each replicate, 30 female mosquitoes were used for each experimental treatment. Data were analyzed by a Log-rank (Mantel-Cox) test using GraphPad Prism 6.0.

### Phagocyte depletion impairs “early-” and “late-phase” mosquito immunity

Several studies have implicated the important role of hemocytes in anti-*Plasmodium* immunity (10, 12, 15, 18, 20, 28, 43), yet the specific contributions of phagocytic immune cells in these immune responses have remained elusive. To investigate the specific role of phagocytes on malaria parasite survival, control and CLD-treated mosquitoes were challenged with *P. berghei* infection (Fig. 3). Phagocyte depletion significantly increased mature oocyst numbers at day 10 (Fig. 3A), indicating that phagocytes serve as critical determinants of parasite survival. When further examined temporally, early oocyst numbers were significantly increased 2 days post-infection (Fig. 3B). With integral roles in ookinete lysis (33, 50, 51), we examined TEP1 expression in mosquito hemolymph in control and clodronate-treated mosquitoes (Fig. 3C). TEP1 protein levels in naïve or *P. berghei*-infected mosquito hemolymph were not influenced by phagocyte depletion (Fig. 3C), however in CLD-treated mosquitoes, TEP1 binding to invading ookinetes was significantly impaired (Fig. 3D). These data argue that mosquito phagocytes mediate TEP1 recognition of invading ookinetes, and is supported by recent findings arguing that hemocyte-derived microvesicles deliver critical factors needed to establish mosquito complement binding to the parasite surface (28).

**Figure 3.**
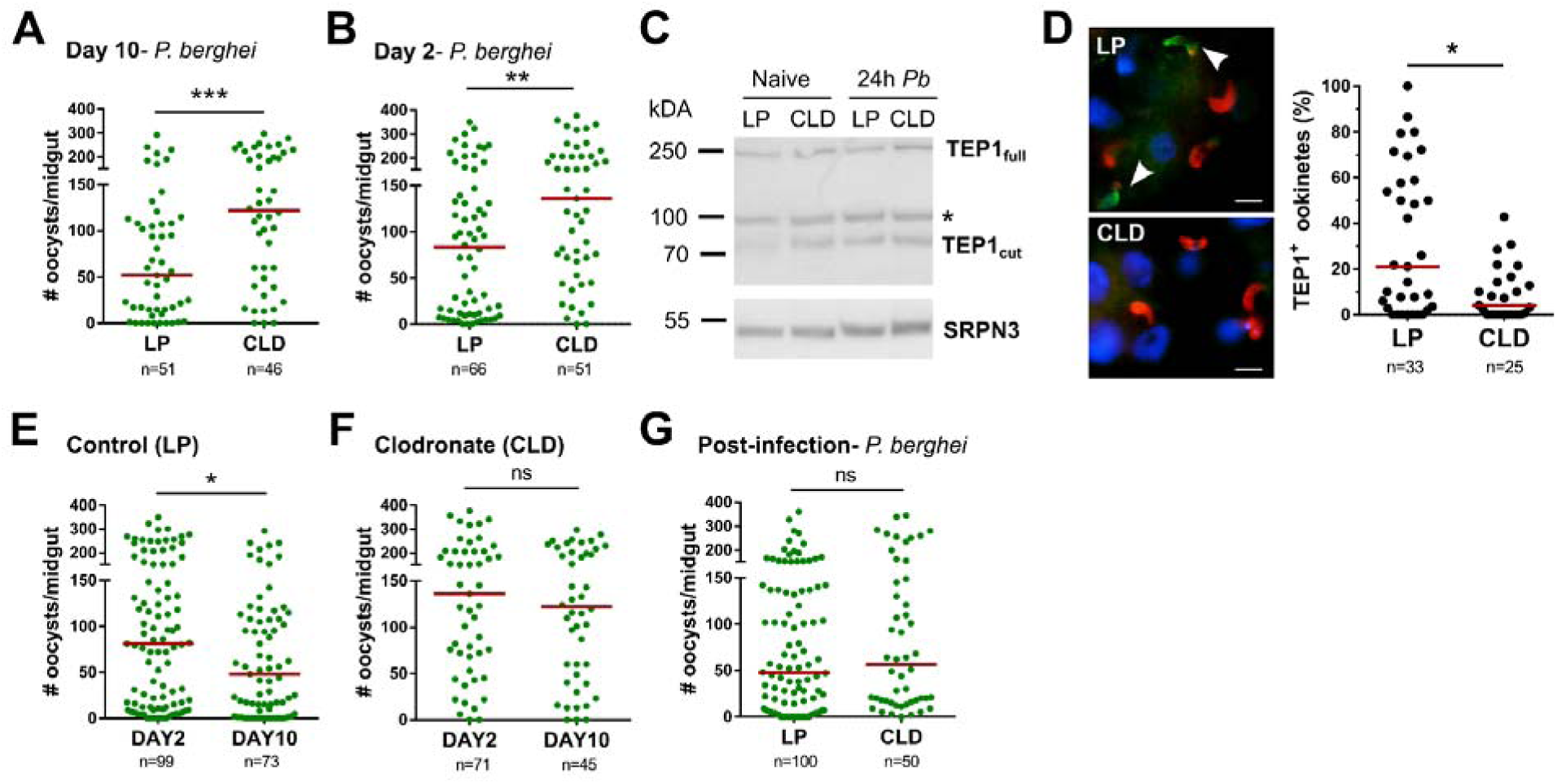
Effects of phagocyte depletion on *P. berghei* development. One day prior to challenge with *P. berghei* (pre-treatment), mosquitoes were treated with control (LP) or clodronate liposomes (CLD). Ten days post-infection, *Plasmodium* oocyst numbers were evaluated to determine the effects of phagocyte depletion on malaria parasite numbers (**A**). To determine the temporal components that influence this increase in parasite survival, day 2 early oocyst numbers were examined in LP- and CLD-treated mosquitoes (**B**). Although phagocyte depletion increased early oocyst numbers, levels of TEP1 protein did not differ between LP and CLD treatments in either naïve or 24 h *P. berghei*-infected hemolymph samples where serpin 3 (SRPN3) was used as a protein loading control (**C**). Non-specific bands were denoted by an asterisk. Evaluation of TEP1 binding to invading ookinetes (~22-24 h post-*P. berghei* infection) by immunofluorescence after phagocyte depletion demonstrates that TEP1 binding (green; indicated by arrows) to ookinetes (a-Pbs 21; red) is significantly impaired (**D**). To examine oocyst survival, oocyst numbers were examined at 2 and 10 days post-*P. berghei* infection in mosquitoes treated with control liposomes (**E**) or clodronate liposomes (**F**). Oocyst numbers were measured by fluorescence using the same cohort of mosquitoes for both time points. Clodronate treatment after the establishment of infection (24 h post-*P. berghei* infection) had no effect on malaria parasite survival (**G**). Three or more independent experiments were performed for all infection experiments and data were analyzed using Mann–Whitney test with GraphPad Prism 6.0. Median oocyst numbers are indicated by the horizontal red line and asterisks denote significance (^⋆^P < 0.05, ^⋆⋆^P < 0.01, ^⋆⋆⋆^P < 0.001); ns, not significant; n, number of midguts examined.

Previous work has also demonstrated the integral role of hemocytes in defining oocyst survival (20, 21). To examine the effects of phagocyte depletion in limiting oocyst numbers, early and late oocyst were measured. In control mosquitoes, oocysts were significantly reduced between Day 2 and Day 8 (Fig. 3E), in agreement with previous descriptions of mosquito late-phase immunity (20–22, 52). However, in CLD-treated mosquitoes, oocyst survival was increased (Fig. 3F), suggesting that phagocytes contribute to both early- and late-phase immune responses that limit parasite numbers in the mosquito host.

Further experiments in which mosquitoes were treated with CLD ~24 h after *P. berghei* challenge demonstrate that there are temporal aspects to the anti-*Plasmodium* immune responses that correspond with phagocyte depletion. In contrast to the results of Fig. 3A, phagocyte depletion after ookinete invasion had no effect on malaria parasite survival (Fig. 3G) although phagocytes were significantly depleted by 77% following CLD treatment after the infection had been established (Fig. S7). Importantly, this suggests that immune responses mediated by phagocytes are induced shortly after ookinete invasion (<24 h), that once these immune mechanisms have been initiated, mosquito phagocytes are no longer required.

### RNA-seq reveals changes in gene expression associated with phagocyte depletion

Evidence form *Drosophila* argues that hemocyte-derived signals are required to initiate humoral immune responses (53, 54), which may similarly contribute to anti-*Plasmodium* immunity in the mosquito host. To better understand the effects of phagocyte depletion on mosquito immunity, RNA-seq analysis was performed on control and CLD-treated mosquito carcass samples 24 h after *P. berghei* infection. To our surprise, only 50 transcripts were differentially regulated (Table S3), of which the majority had annotated immune function (Fig. 4A). This included the known phagocyte proteins LRIM 16A, LRIM 16B, and nimrod B2 (10), as well as several other leucine rich-repeat (*LRR*) proteins, fibrinogens, and multiple prophenoloxidase (*PPO*) genes (*PPO*-*2*, -*3*, -*4*, -*5*, -*6*, -*9*) that were significantly down-regulated following CLD treatment (Fig. 4B). Additional validation of the RNA-seq data was performed using qRT-PCR analyses on select genes, producing comparable levels of gene expression and a significant correlation (R^2^=0.94) between both types of analyses (Fig. S8).

**Figure 4.**
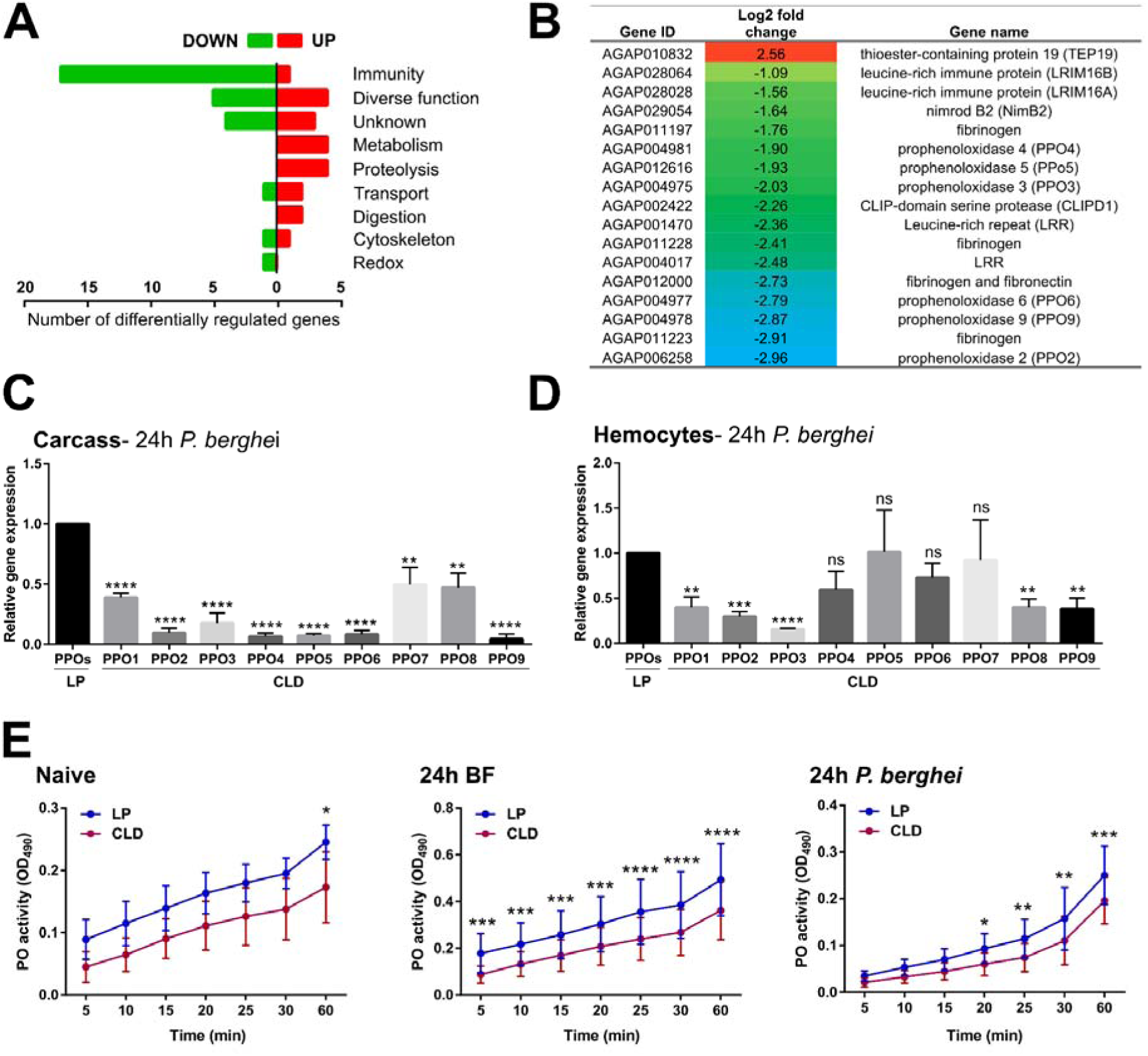
Phagocyte depletion reduces of prophenoloxidase (*PPO*) expression. RNA seq analyses following clodronate treatment revealed 50 differentially regulated genes in abdomen tissues 24 h post-*P. berghei* infection and grouped by gene ontology (**A**). Comprising the largest category of affected genes, the annotations and log2 fold change of specific immune genes with significant differential regulation are displayed (**B**). This includes several PPO genes, therefore leading us to examine the expression of all nine PPO family members by qRT-PCR analyses in the carcass (**C**) and hemocyte (**D**) samples in control liposome (LP) and clodronate-treated (CLD) samples. Data were analyzed using an unpaired f-test to determine differences in relative gene expression of each respective PPO gene between LP and CLD treatments (**C** and **D**). Due to the importance of PPOs in phenoloxidase (PO) activation, PO activity was measured in hemolymph samples derived from LP and CLD samples in naïve, blood-fed, and *P. berghei*-infected conditions (**E**). Measurements (OD_490_) were taken for DOPA conversion assays at five minute intervals from 0 to 30 minutes, then again using a final readout at 60 min. Data were analyzed using a two-way repeated measures ANOVA followed by Sidak’s multiple comparisons using GraphPad Prism 6.0. Bars represent mean ± SEM of three independent experiments. Asterisks denote significance (^⋆^*P* < 0.05, ^⋆⋆^*P* < 0.01, ^⋆⋆⋆^*P* < 0.001, ^⋆⋆⋆⋆^*P* < 0.0001).

### PPO expression and phenoloxidase activity are significantly reduced following phagocyte depletion

With the identification of multiple PPO genes influenced by phagocyte depletion (Fig. 4B), we further explored this conspicuous result by examining the expression of all 9 annotated PPO genes. When *PPO* expression was examined in the carcass, all 9 *PPO* genes displayed a significant reduction in their expression (Fig.4C), similar to our RNA-seq analysis. In contrast, when perfused hemocytes were examined (Fig. 4D), only *PPO*-*2*, -*3*, -*8*, -*9* expression were significantly reduced following CLD-treatment arguing that some *PPOs* are still expressed in circulating hemocytes. Serving as precursors of melanization reactions in insects (55), phenoloxidase (PO) activity was measured in perfused hemolymph from control and CLD-treated mosquitoes. 48 h post-treatment, PO activity was reduced in naïve CLD treated mosquitoes displaying significant differences after 60 min (Fig. 4E). The influence of blood feeding amplified these responses, reducing PO activity across all sample time points (Fig. 4E). Similarly, PO activity was reduced following *P. berghei* infection, although the kinetics of the reaction were slower and produced lower levels of activity by comparison to the non-infected blood feeding (Fig. 4E). Together, these data imply that *PPO* expression and subsequent PO activity are impaired following phagocyte depletion.

### Multiple PPOs influence *Plasmodium* oocyst survival

Based on the RNA-seq results and reduced PO activity in CLD-treated mosquitoes (Fig. 4), we examined several candidate immune genes that featured prominently in our analyses to assess their respective contributions to *Plasmodium* survival using RNAi. For this reason, we examined *CLIPD1*, a putative leucine-rich immune protein (LRIM) AGAP001470, as well as multiple PPO genes (*PPO*-*2*, -*3*, -*4*, -*5*, -*6* and -*9*). All eight candidate genes were significantly silenced following the injection of dsRNA (Fig. S9). As members of a multi-gene family, the specificity of *PPO* silencing was further verified using specific primers for each of the nine PPO genes in *An. gambiae* (Fig. S10). Gene-specific dsRNA constructs specifically targeted *PPO2*, *PPO4*, *PPO5*, and *PPO6*, while *PPO3* and *PPO9* had unintended off-target effects on one or more PPO genes (Fig. S10). The loss of *CLIPD1*, *AGAP001470*, *PPO4*, *PPO5*, and *PPO6* had no effect on parasite numbers (Fig. S11), while silencing *PPO2*, *PPO3*, or *PPO9* increased the intensity of malaria parasite infection at day 8 (Fig. 5A). However, when early oocyst numbers were examined at day 2, *PPO2*-, *PPO3*-, and *PPO9*-silencing did not influence ookinete invasion success (Fig. 5B), suggesting that *PPO2*, *PPO3*, and *PPO9* contribute to oocyst survival. Similar to previous reports (20, 21, 52), oocyst numbers were compared at day 2 and day 8 from the same cohort mosquitoes after silencing *PPO2.* Oocyst numbers significant decrease in control GFP mosquitoes between day 2 and day 8, while these effects are abrogated in *PPO2* silenced mosquitoes (Fig. 5C), together suggesting that *PPOs* are important determinants of oocyst survival.

**Figure 5.**
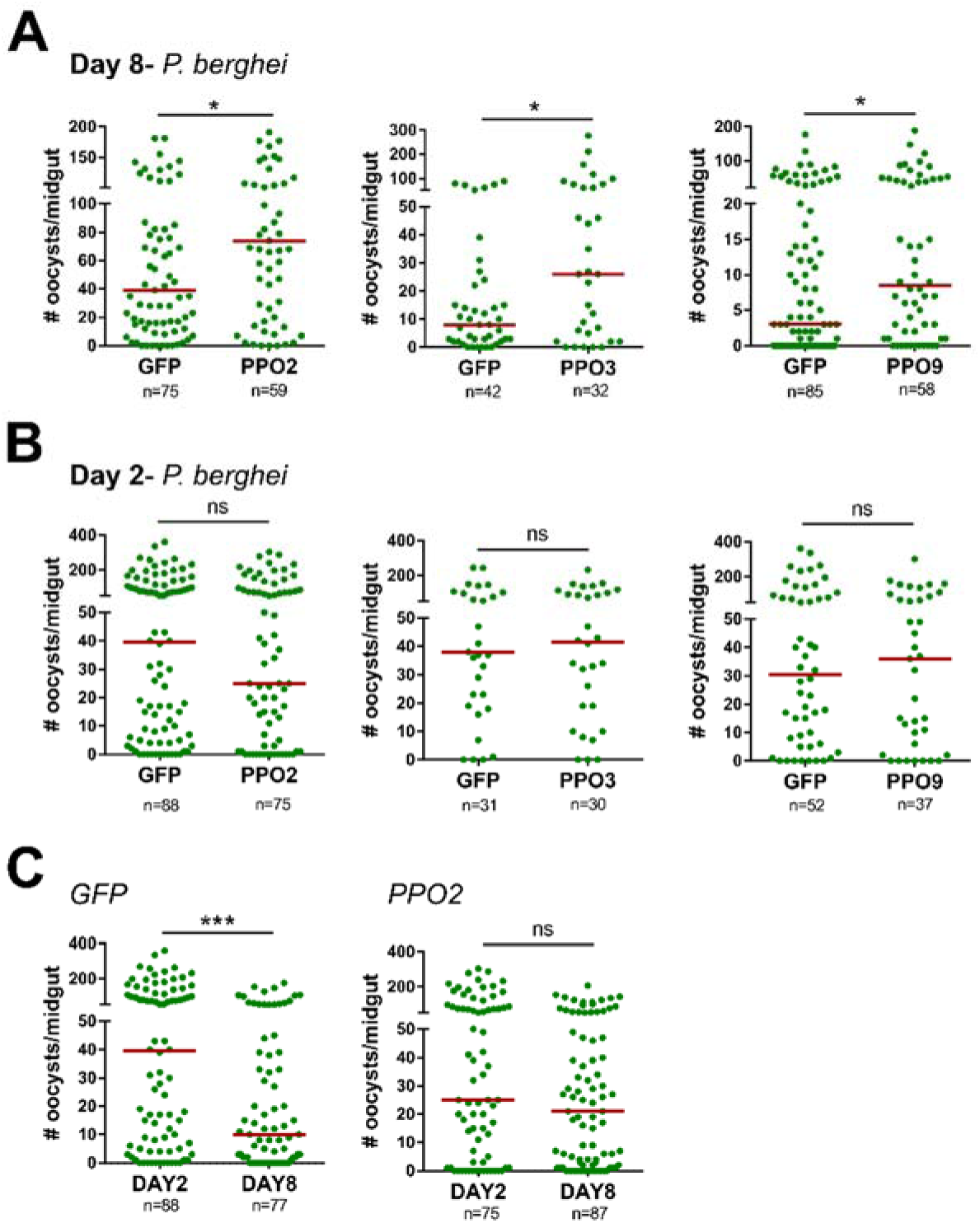
The silencing of PPOs influences *Plasmodium* oocyst survival. The influence of *PPO2*-, *PPO3*-, and *PPO9*-silencing on *Plasmodium* oocysts numbers in *An. gambiae* was evaluated 8 days post-infection as compared to dsGFP controls (**A**). To determine if gene-silencing influences the success of ookinete invasion, similar experiments were performed in which early oocyst numbers were used as a readout of ookinete survival two days post-infection (**B**). *PPO2*-silenced mosquitoes were further examined at 2 and 8 days post-infection with *P. berghei* to validate the contributions of PPO2 on oocyst survival (**C**). Data were collected using the same cohort of mosquitoes for both time points. For all experiments, each dot represents the number of parasites on an individual midgut, with the median value denoted by a horizontal red line. Data were pooled from three or more independent experiments with statistical analysis determined by a Mann–Whitney test using GraphPad Prism 6.0. Asterisks denote significance (^⋆^*P* < 0.05, ^⋆⋆⋆^*P* < 0.001).

### Analyses of PPO6 protein expression in phagocytes

While PPO6-silencing does not produce a discernable oocyst phenotype, several reports have described PPO6 as a reliable marker of mosquito hemocytes (8, 9, 56, 57) and phagocytes (10, 57). Following CLD-treatment, the number of circulating PPO6^+^ phagocytes are significantly depleted (Fig. 6A), resulting in a higher proportion of non-phagocytic PPO6^+^ cells comprising the remaining immune cell population (Fig. 6B). This provides support that PPO6^+^ phagocytes are depleted by CLD-treatment, yet hemolymph expression of PPO6 remains unchanged (Fig. S12), suggesting that additional immune cell sub-types without phagocytic properties, possibly oenocytoids, may contribute to its production.

**Figure 6.**
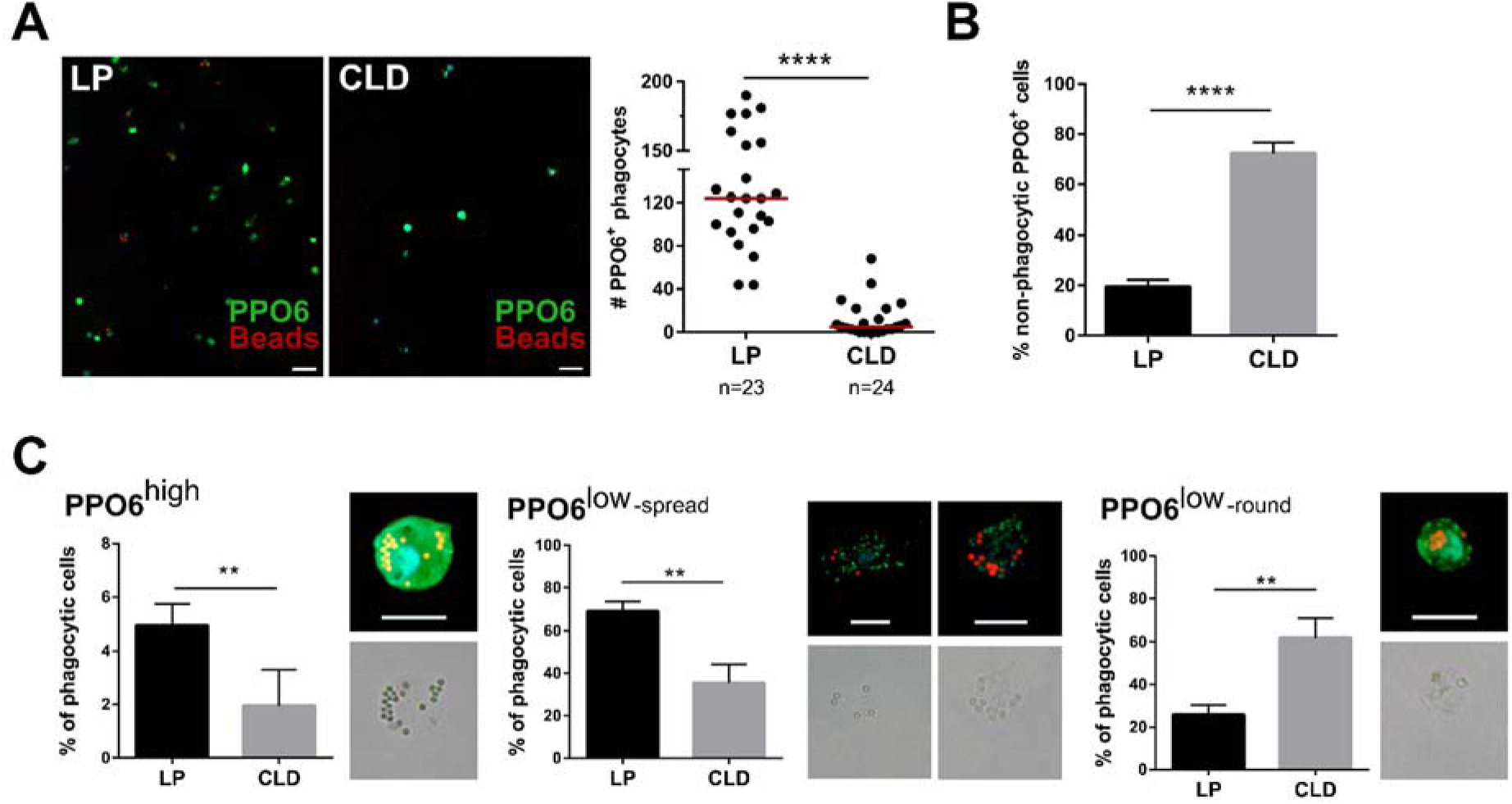
Clodronate treatment differentially impacts mosquito phagocyte subpopulations. PPO6 staining was evaluated by immunofluorescence in perfused hemocytes after the injection of fluorescent beads from control liposome (LP) or clodronate-treated (CLD) mosquitoes ~24hr post-infection with *P. berghei.* After adherence and fixation, hemocytes were stained with a PPO6 antibody (**A**). PPO6^+^ phagocytes (green) that have phagocytosed red fluorescent beads were compared between LP and CLD treatments, as well as the proportion of non-phagocytic PPO6^+^ cells (**B**). Closer examination of PPO6^+^ phagocytic cells revealed distinct populations of immune cells distinguished by PPO6 signal intensity (high or low) and morphological features (elongated/spread, small rounded) (**C**). Two independent experiments of immunofluorescent assays were performed. Data were analyzed by Mann–Whitney test using GraphPad Prism 6.0. Median is indicated by the horizontal red line (**A**). Bars represent mean ± SEM (**B** and **C**). Asterisks denote significance (^⋆⋆^*P* < 0.01, ^⋆⋆⋆⋆^*p* < 0.0001). Scale bar: 10 μm.

In addition, similar to Severo et al. (57), we identify variable (high and low) expression of PPO6 expression in PPO6^+^ immune cells with and without phagocytic ability (Fig. S13). Based on these high or low PPO6 expression phenotypes, additional spreading/elongated or rounded morphological features were used to further define these cell populations following phagocyte depletion (Fig. 6C). CLD-treatment was most effective in reducing populations of PPO6^low^ cells with elongated or spread morphologies most commonly associated with mosquito granulocytes (Fig. 6C). Phagocytic PPO6^high^ cell populations also displayed significantly reduced cell proportions following CLD-treatment, although these cell types comprise a much smaller proportion of phagocytic cells (Fig. 6C). Interestingly, a small, rounded PPO6^low^ phagocytic cell similar to those described by King and Hillyer (6), comprised the majority of phagocytic cells following CLD-treatment and significantly differed from control mosquitoes (Fig. 6C). Based on these results, this small PPO6^low^ phagocytic cell population may be less susceptible to CLD-treatment and likely comprise the “lower” cell population described in Fig. S6. Together, these data provide further support that clodronate liposomes can specifically deplete distinct sub-populations of phagocytic immune cells in the mosquito host, and provide new insight into the complexity of mosquito phagocyte populations.

## Discussion

Over the last decade, we have gained significant insights into the biology of mosquito hemocytes and their respective roles in innate immune function. From gene and protein expression studies (10, 12, 13), an extensive catalog of the molecular responses to blood feeding and pathogen challenge has been produced. New information into the impacts of physiology on mosquito immune cell dynamics (7–9), infection-induced interactions of circulating hemocytes (6, 11, 58), and the role of hemocytes as important modulators of anti-*Plasmodium* immunity (10, 12, 16, 18, 20, 28, 43) have recently been addressed. However, the functional classifications of these mosquito immune cell populations has been severely limited by the lack of genetic tools and molecular markers.

Relying primarily on morphological identification, mosquito hemocytes have been broadly characterized as prohemocytes, oenocytoids, and granulocytes based on homology to other invertebrate systems (3, 4). Of these cell types, granulocytes have been the most well-studied for their distinct shape and phagocytic properties, resulting in the identification of several immune molecules with integral roles in phagocytosis (14, 15) and enabling a phagocyte-specific proteome (10). Similarly taking advantage of these phagocytic properties, in this study we describe the ability to chemically deplete *An. gambiae* phagocytes through the use of clodronate liposomes.

Widely used in mammalian systems, clodronate liposomes have been used to effectively deplete vertebrate macrophage cell populations (23–25). Following phagocytosis by phagocytic immune cells, liposome particles are believed to be broken down by the endosome, thus releasing clodronate that promotes apoptosis and the depletion of phagocytic cells (23, 24). Used for the first time in an invertebrate system herein, we demonstrate that clodronate liposomes can effectively deplete phagocytic cell populations in the malaria vector, *An. gambiae*, through the validation of a variety of cellular- and molecular-based approaches. Light microscopy and immunofluorescence assays argue that granulocyte populations are significantly reduced following CLD-treatment, which is further supported by the decrease in relative transcript levels of *eater* and *nimrod B2* that are routinely associated with phagocytic hemocytes in mosquitoes (47, 48) and other Diptera (45, 46). Additional validation by flow cytometry analysis confirmed the effects of clodronate treatment on phagocyte depletion, as well as revealed at least two different phagocytic cell populations in mosquitoes with varied levels of susceptibility to clodronate treatment. When combined with the presence/absence of PPO6 staining following CLD treatment, our data suggest that three or more phagocyte populations are present in *An. gambiae.* This consists of highly phagocytic PPO6^+^ and PPO6^-^ cell types, similar to those described by Severo et al. (57), that are susceptible to phagocyte depletion. An additional small, rounded PPO6^-^ phagocytic cell similar to those described by King and Hillyer (6), displayed little response to CLD treatment. Together, these data provide strong evidence that clodronate liposomes can be used to deplete mosquito phagocytic cell populations, while providing new insights into previously undescribed complexities of mosquito hemocyte populations.

From these experiments, we provide evidence that the efficacy of clodronate treatment is influenced by mosquito physiology. Phagocyte depletion was much more effective following blood-feeding, independent of pathogen challenge. While the mechanisms for this increased phagocytic activity remain unknown, other studies have reported increased cellular activity and up-regulation of immune related molecules in hemocytes following blood feeding (9, 10), which may lead to an increased capacity to ingest clodronate liposomes under these physiological conditions.

Infection experiments following phagocyte depletion demonstrate the integral role of phagocytic immune cells in the mosquito immune response to bacterial and parasitic pathogens. With phagocytosis serving as an integral mechanism to remove invading pathogens, it is to some extent not surprising that mosquito survival is significantly reduced upon bacterial challenge following clodronate treatment. Challenge experiments with both gram (+) and gram (-) bacteria caused significant mortality in mosquitoes, as early as one day post-challenge with *S. marcesens.* Demonstrating the vital role of phagocytes on host survival, these data provide additional functional validation of the ability to deplete phagocytic immune cells in the mosquito host.

With the use of clodronate liposomes to deplete phagocyte populations, these novel experiments have enabled the ability to address the specific contributions of phagocytic immune cell sub-types to malaria parasite infection for the first time. Previous work implicating mosquito hemocytes and hemocyte-derived components have been limited to reverse genetic approaches evaluating the influence of candidate gene function (10, 12, 16, 18, 20, 28) or in over-loading the phagocytic capacity of cells to evaluate their cellular function (28, 43). Through our experiments using clodronate liposomes, we now provide definitive insight into the roles of phagocytic immune cells to anti-*Plasmodium* responses in the mosquito host. These data suggest that mosquito phagocytes mediate multi-modal immune responses that target both the ookinete and oocyst stages of malaria parasite development through distinct immune mechanisms as previously proposed (20–22).

Recognition and lysis of invading ookinetes occurs at the interphase of the basal lamina, where malaria parasite first become exposed to components of the mosquito hemolymph (22, 59). There mosquito complement recognition directs ookinete killing responses that require TEP1 function (33, 50, 51). Following phagocyte depletion, early oocyst numbers are significantly increased, suggesting that clodronate treatment increases the survival of invading ookinetes. As a result, we therefore examined the effects of phagocyte depletion on TEP1 expression. Although circulating levels of TEP1 were not altered in mosquito hemolymph following CLD treatment, TEP1 binding to invading ookinetes was significantly impaired. This is in agreement with recent studies by Castillo et al. (28) that argue that hemocyte-derived microvesicles (HdMV) are critical mediators of TEP1 binding to invading ookinetes, suggesting that phagocytic immune cells produce the HdMV required for ookinete lysis.

In addition to these “early-phase” immune responses, phagocyte depletion via clodronate liposomes also increased *Plasmodium* oocyst survival similar to previously characterized “late-phase” phenotypes (20–22, 52). Previous results have implicated hemocytes in this process (20, 21), yet through the use of clodronate liposomes we can confirm the specific involvement of phagocytes in mediating oocyst killing responses. Importantly, our results shed new insight into the mechanisms of late-phase immunity through the identification of multiple PPOs that are dysregulated following phagocyte depletion. Through gene-silencing experiments, we demonstrate the ability to selectively target individual PPO genes to evaluate the contributions of six individual PPOs on malaria parasite survival. From these experiments, we identify 3 PPO genes, PPO2, PPO3, and PPO9 that limit the survival of *Plasmodium* oocysts. Most commonly associated with melanization, the role of PPOs in oocyst killing has not been fully elucidated, but the lack of melanized oocysts in our experiments suggest a different mechanism of action. In addition to the production of melanin, PPOs have been implicated in coagulation and wound healing responses that contribute to the elimination of bacterial, viral, and parasitic pathogens (60–64). As a result, we believe that these killing responses are likely mediated by cytotoxic intermediates produced by the activation of the phenoloxidase (PO) cascade (65). This includes reactive oxygen or nitrogen intermediates that may be permeable to the midgut basal lamina that otherwise protects maturing oocysts from components of the mosquito hemolymph. However, the exact mechanisms by which a subset of PPOs promote parasite killing have yet to be identified and are the focus of future work.

While the role of PPOs in innate immunity has been well studied in other insects (60, 66), our current understanding of PPOs in mosquitoes has been limited and further complicated by their recent gene expansion in mosquito species. *An. gambiae* has nine annotated PPO genes, compared to only three identified in *Drosophila.* Based primarily on evidence from other insect systems, mosquito oenocytoid populations have predominantly been implicated in the expression of mosquito PPOs. However, recent studies have begun to illustrate that phagocytic cell populations are also important components of mosquito PPO production (10, 57), in agreement with our results following phagocyte depletion. This is further supported by additional studies examining PPO6 transgene expression in *An. gambiae* (56, 57), and PPO staining in granulocytes of mosquitoes, moths, and houseflies (2, 8, 9, 67, 68) that together argue that phagocytic cell populations have integral roles in PPO expression and PO activity. Moreover, we believe that these functional characterizations and differences in PPO expression make a case to revisit the morphological classifications of mosquito hemocyte populations from prohemocytes, oenocytoid, and granulocytes into molecular classifications that are more indicative of cellular function. Following this methodology, our data and that of Severo et al (57), point to at least three phagocytic cell populations that can be defined by PPO6 expression and morphological characteristics. Therefore, these insights extend well beyond the “granulocyte” classification that have previously denoted mosquito phagocytic cell populations and provide new details into the complexity of these distinct cell populations.

An additional, interesting consideration of our work, are the temporal aspects of cellular immunity in malaria parasite killing. While clodronate treatment and subsequent phagocyte depletion prior to infection lead to significant increases in ookinete and oocyst survival, treatment after an infection had been established (~24 h post-infection) had no effect on malaria parasite numbers. This suggests that the presence of phagocytic immune cell populations during the time of ookinete invasion are able to initiate both “early-” and “late-phase” immune responses respectively targeting *Plasmodium* ookinetes and oocysts (Fig. 7), that when ablated significantly increase parasite survival. However, when phagocytes are depleted by clodronate treatment after ookinete invasion has occurred, the immune signals that contribute to these immune killing responses have already been set in motion. This is in agreement with previous models of early- and late-phase immunity that suggest that these immune signals are initiated in response to midgut epithelial damage (20, 22), where parasites are likely killed through non-specific to cellular damage and wound healing.

**Figure 7.**
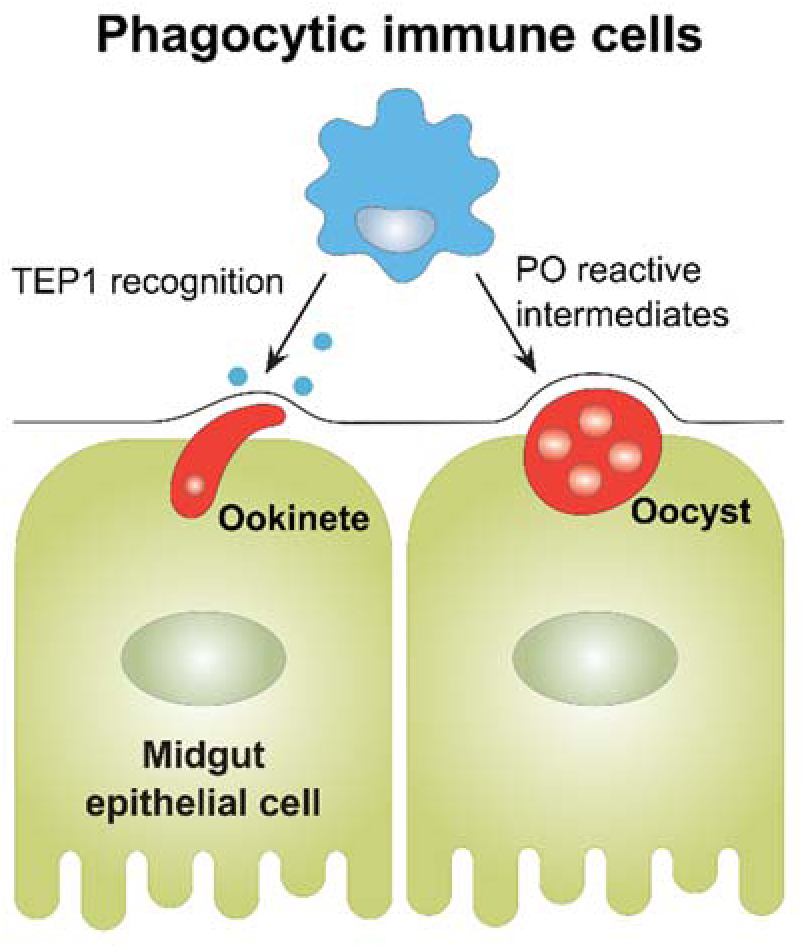
Multimodal contributions of phagocytes on anti-*Plasmodium* immunity. Experiments with clodronate liposomes establish integral roles of phagocytic immune cells in malaria parasite killing which include the role of phagocytes in the recognition of invading ookinetes and the production of prophenoloxidases that limit oocyst survival.

In summary, our findings are the first definitive characterization of mosquito phagocytes in anti-*Plasmodium* immunity. Taking advantage of tools from vertebrate systems, we demonstrate that clodronate liposomes are an effective chemical method to deplete phagocytic immune cells in *An. gambiae*, thus enabling methodologies to better understand the immune contributions of phagocytic immune cells and malaria parasites in the mosquito host. From these data, we corroborate recent studies arguing for the role of cellular immunity in directing the recognition and killing of *Plasmodium* ookinetes, as well as provide new insights into the mechanisms that influence oocyst survival. These studies also shed important new information into the complexity of phagocytic cell populations, identifying at least three types of phagocytic immune cells, highlighting the need for further molecular characterization of immune cell populations in invertebrate systems. Together, we believe our study represents a significant advancement in our understanding of mosquito immune cells and the mechanisms by which they contribute to malaria parasite killing.

## Acknowledgments

The authors would like to thank Andrea Radtke for the initial discussions that led this project. This work would not have been possible without the generosity of Kristin Michel for sharing the SRPN3 antibody, Michael Povelones for providing the TEP1 antibody, and to George Christophides for offering the PPO6 antibody used in these studies. Additional help was provided by Linda Zeller for providing the *S. marcescens* and *S. aureus* bacterial cultures used in bacterial challenge experiments. Special thanks to Shaun Rigby of the Iowa State Flow Cytometry Facility for his instrumental help in performing flow cytometry experiments. We also thank Arun Somwarpet-Seetharam and Andrew Severin of the Iowa State Genome Informatics Facility for assistance with the gene expression data, Mike Baker of the Iowa State DNA Core Facility for assistance with the RNA-seq project, and Hee Jung Oh for assistance in scientific illustrations. This research was supported in part by the Agricultural Experiment Station at Iowa State University.

